# TatA complexes exhibit a marked change in organisation in response to expression of the TatBC complex

**DOI:** 10.1101/079715

**Authors:** Sarah M. Smith, Andrew Yarwood, Roland A. Fleck, Colin Robinson, Corinne J. Smith

## Abstract

The twin arginine translocation (Tat) system is an integral membrane protein complex that accomplishes the remarkable feat of transporting large, fully-folded polypeptides across the inner membrane of bacteria, into the periplasm. In *Escherichia coli* Tat is comprised of three membrane proteins: TatA, TatB and TatC. How these proteins arrange themselves in the inner membrane to permit passage of Tat substrates, whilst maintaining membrane integrity, is still poorly understood. TatA is the most abundant component of this complex and facilitates assembly of the transport mechanism. We have utilised immunogold labelling in combination with array tomography to gain insight into the localisation and distribution of the TatA protein in *E. coli* cells. We show that TatA exhibits a uniform distribution throughout the inner membrane of *E. coli* and that altering the expression of TatBC shows a previously uncharacterised distribution of TatA in the inner membrane. Array tomography was used to provide our first insight into this altered distribution of TatA in 3D space, revealing that this protein forms linear clusters in the inner membrane of *E.* coli upon increased expression of TatBC. This is the first indication that TatA organisation in the inner membrane alters in response to changes in Tat subunit stoichiometry.

**Summary statement:** The volumetric electron-microscopy technique, array tomography, revealed a novel distribution of TatA protein (from the twin arginine translocase complex), in *Escherichia coli*.

## Introduction

The bacterial Tat system transports fully-folded proteins across the bacterial inner membrane and the chloroplast thylakoid membrane to the periplasm (reviewed by(Patel, 2014, Cline, 2015)). Tat is important for the targeting of proteins involved in numerous important processes including energy metabolism, formation of the cell envelope and nutrient acquisition. There is also evidence that Tat is required in bacterial pathogenesis (Ochsner et al., 2002, McDonough et al., 2008). In Gramnegative bacteria the Tat system is comprised of three membrane proteins: TatA, TatB and TatC (Bogsch et al., 1998, Weiner et al., 1998). Substrates are targeted to the Tat machinery *via* an N-terminal signal peptide, which contains an invariant twin-arginine motif (Chaddock et al., 1995, Stanley et al., 2000). At the target membrane the substrate makes contact with a TatBC complex containing multiple copies of these subunits (Tarry et al., 2009, Ma and Cline, 2013). Interaction with this substrate-binding complex triggers the recruitment of a separate TatA complex (Alami et al., 2003, Mori and Cline, 2002) to form the active translocase through which the substrate can pass. The precise mechanism of translocation is, however, poorly understood and the full translocation system has not been characterised.

It has been proposed that TatA contributes to the translocation process either by weakening the membrane, or by forming a transient pore. Negative stain electron microscopy structures of detergent-solubilised *E. coli* TatA complexes showed them to be present as large, homo-oligomeric complexes that possess a cavity of varying diameter (Gohlke et al., 2005). These data provided support for the idea that TatA contributes the pore-forming element of the Tat machinery. However, similar studies on *E. coli* TatE (a TatA paralogue that can substitute for TatA) and *Bacillus subtilis* TatAd (which can also substitute for TatA) showed these complexes to be present as much smaller, fairly homogeneous particles (Baglieri et al., 2012, Beck, 2013). The true significance of Tat heterogeneity thus remains unclear.

Alternatively, it has been proposed that once TatA is recruited to the TatBC complex it could serve as a nucleation point for additional TatA proteins (Alami et al., 2003, Mori and Cline, 2002). Cross-linking data has shown that the number of cross-links between TatA proteins increases under transport conditions (Dabney-Smith et al., 2006), which is consistent with the idea that substrate binding to the Tat machinery triggers polymerisation of TatA. This in turn would alter the immediate environment of the phospholipid bilayer to permit passage of substrate through the inner membrane (Dabney-Smith et al., 2006, Brüser and Sanders, 2003, Cline and McCaffery, 2007, Greene et al., 2007). However, the precise oligomeric state of the TatA protein varies in cell membranes and detergent micelles (Gohlke et al., 2005, Porcelli et al., 2002, Leake et al., 2008, Alcock et al., 2013), and the net result is that the true significance of TatA oligomerisation remains unclear.

The distribution of Tat complexes within the membrane is another area that has been difficult to analyse with accuracy. Efforts to investigate the distribution of the two Tat complexes (TatA and TatBC) have utilised Tat proteins tagged with fluorescent proteins, and there is evidence that the TatB and TatC subunits retain activity when C-terminal fluorescent tags are present (Ray et al., 2005). This study concluded that these proteins were uniformly distributed when expressed at low levels, although a separate study proposed that Tat complexes adopted a polar localisation (Berthelmann and Brüser, 2004). However, analysis of TatA is more difficult owing to the fact that studies which used fluorescently-tagged TatA have produced ambiguous results: one study used GFP-tagged TatA to analyse the oligomeric state of TatA complexes in the membrane (Leake et al., 2008), but another (Ray et al., 2005) showed that a TatA-GFP fusion is prone to proteolysis and release of mature-size TatA. Since very low levels of TatA can support translocation activity, this finding means that the use of TatA-GFP fusion proteins to measure TatA protein distribution and/or localisation in two dimensions is problematic.

Analysis of the 3-dimensional distribution of Tat complexes within cells is also challenging. Despite the advent of super-resolution light microscopy bringing opportunity to analyse bacterial protein distribution with high resolution(Bakshi et al., 2012, English et al., 2011, Greenfield et al., 2009), it has been hard to visualise Tat protein distribution in 3D space, owed largely to the fact that they are small proteins which only transiently interact at the membrane (Cline and McCaffery, 2007). Coupled with the above questions regarding TatA-GFP activity, this means that care must be taken when considering light microscopy-based approaches to study Tat localisation.

In this study we have sought to overcome such problems through the use of electron microscopic approaches in which native, unmodified TatA proteins are localised directly for the first time. We have developed a novel combination of immunogold labelling and array tomography (Micheva. K.D., 2007) of *E. coli* to unveil the distribution of TatA in the inner membrane (Fig. S1). 2D analysis of immunolabelled *E. coli* shows that TatA is predominantly uniformly distributed in the inner membrane (whether TatA is overexpressed or expressed alongside TatBC). However, a proportion of TatA exhibits a linear clustering along the inner membrane in a TatBC-dependent manner. Array tomography successfully gained preliminary insight into this previously uncharacterised organisation of TatA in the inner membrane, confirming that this protein has the ability to exhibit a higher order of organisation in the cell wall.

Not only is this the first time that *E. coli* has been analysed using array tomography, it is the first time that the localisation and distribution of any Tat component has been visualised in 3D space in its native, cellular environment. This methodology has far-reaching potential for use with any suitable cellular antigen in diverse biological systems.

## Results

### TatA exhibits a uniform distribution in the inner membrane of E. coli in cells overexpressing only TatA

The distribution of TatA was examined in wild type MC100 cells over-expressing TatA (only), and in cells overexpressing TatABC from a plasmid-borne *tatABC* operon. This approach was taken so that we could test whether the presence of stoichiometric levels of the TatBC complex affects the distribution of TatA complexes. *E. coli* cells lacking any Tat machinery (∆*tatABCDE*; hereafter denoted ∆*tat*) were used as controls for non-specific binding.

Cells overexpressing either TatA or TatABC were harvested, normalised (to an OD_600_ of 10.0) and fractionated into cytoplasmic and membrane fractions to determine the proportion of TatA in these cellular fractions (see Figure 1). With cells over-expressing either TatA or TatABC, the vast majority of the TatA is found in the membrane fraction as expected, while a small proportion (about 10-20%) is found in the cytoplasm, as observed in previous studies (Berthelmann et al., 2008). In the control tests, no signal is obtained in blots of ∆*tat* cells that lack the TatA protein.

**Figure 1.**
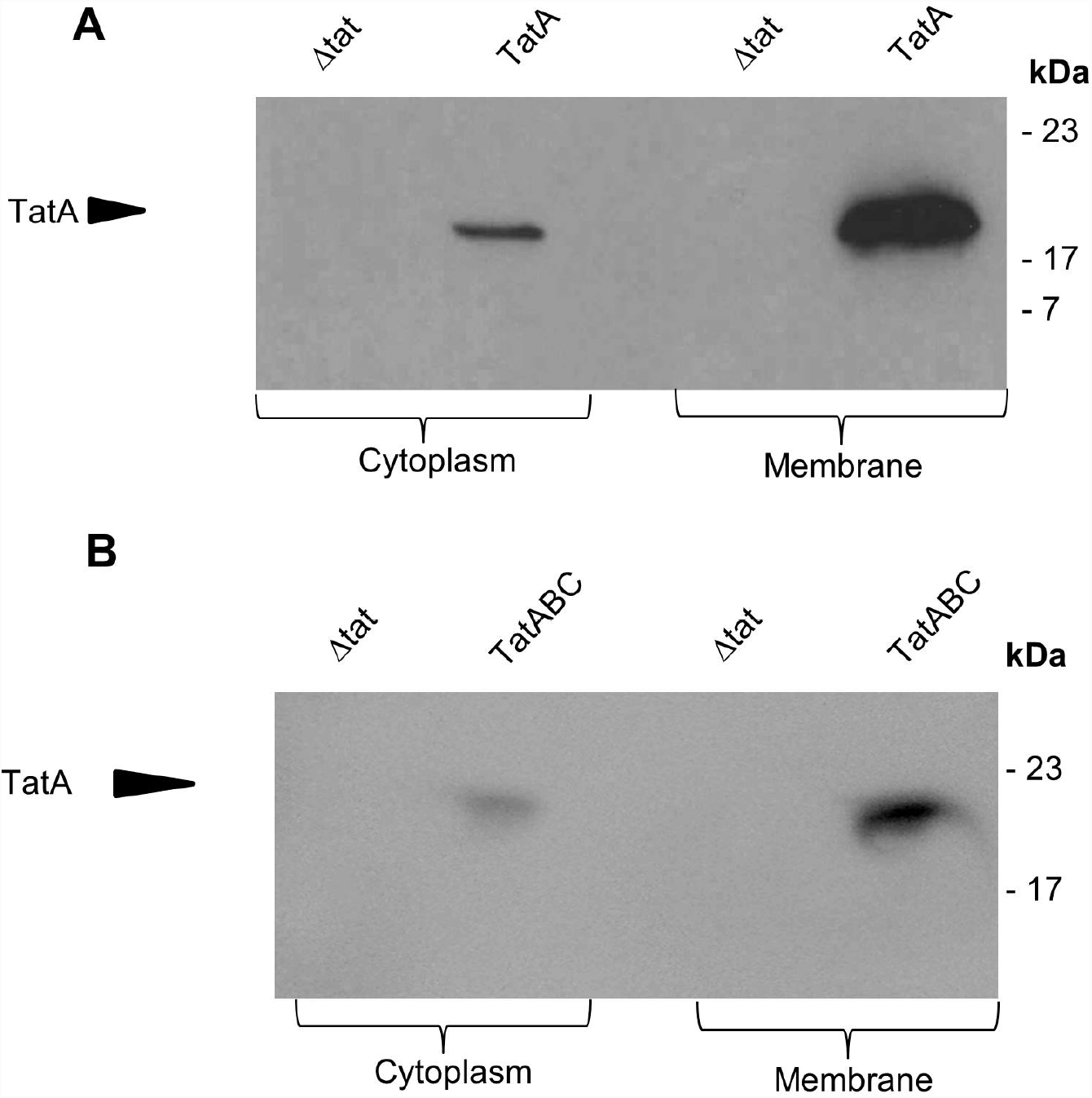
Cytoplasmic and membrane fractions of *E. coli* cells (overexpressing TatABC and TatA) immunoblotted for the detection of TatA protein using anti-TatA antibody. *E. coli* cells overexpressing TatA (A) and TatABC (B) were fractionated into cytoplasmic and membrane fractions. The fractions were resolved by SDS PAGE and immunoblotted to detect for the presence of TatA protein. The control was *E. coli* cells that have had Tat machinery knocked-out (∆*tat*). TatA protein was identified in the cytoplasm and membrane compartment of both cell expressing TatA and TatABC.

We first analysed the distribution of TatA in 2D sections using immunogold labelling. *E. coli* cells overexpressing TatA or TatABC were fixed and stained to preserve the cells and provide contrast; the images were examined for the presence of gold particles and these were classified according to their location. For this study, a gold particle found within 25 nm of the inner membrane was defined as being located at the inner membrane (dimensions are illustrated in Fig. S2). Our aim was to achieve unambiguous identification of TatA protein in the inner membrane through immunogold labelling of *E. coli* cells overexpressing only TatA. In order to do this, the level of non-specific antibody labelling was assessed through comparison of TatA-overexpressing cells with ∆*tat* cells that lack Tat machinery.

Figure 2A shows representative images of TatA-overexpressing cells that were immunogold-labelled with anti-TatA. These cells exhibit a pattern of gold labelling that is predominantly around the periphery of cell. This was confirmed by magnification of the cells, as in Fig. S2, and showed the gold particles to reside within 25 nm of the inner of the cell wall; thus we therefore conclude that these particles were almost entirely located in the inner membrane. A smaller number of particles are evident in the cell interior. Analysis of ∆*tat* cells (representative images shown in Fig. S3) shows the presence of a small number of gold particles in the cytoplasmic region with very few in the membranes. These data confirm that the antibody is able to detect TatA with a high degree of specificity. As further controls, TatA-expressing cells were immunolabelled with the primary antibody omitted (Fig. S4), and TEM analysis showed that these cells lacked any gold binding; this confirms that non-specific binding seen in ∆*tat* cells was attributable to the primary antibody. Finally, the same cell type was immunolabelled with a gold-conjugated secondary antibody directed towards a different animal species (Fig. S3); TEM analysis showed only unlabelled *E. coli* cells, confirming that the cells do not have a non-specific attraction for gold particles.

**Figure 2.**
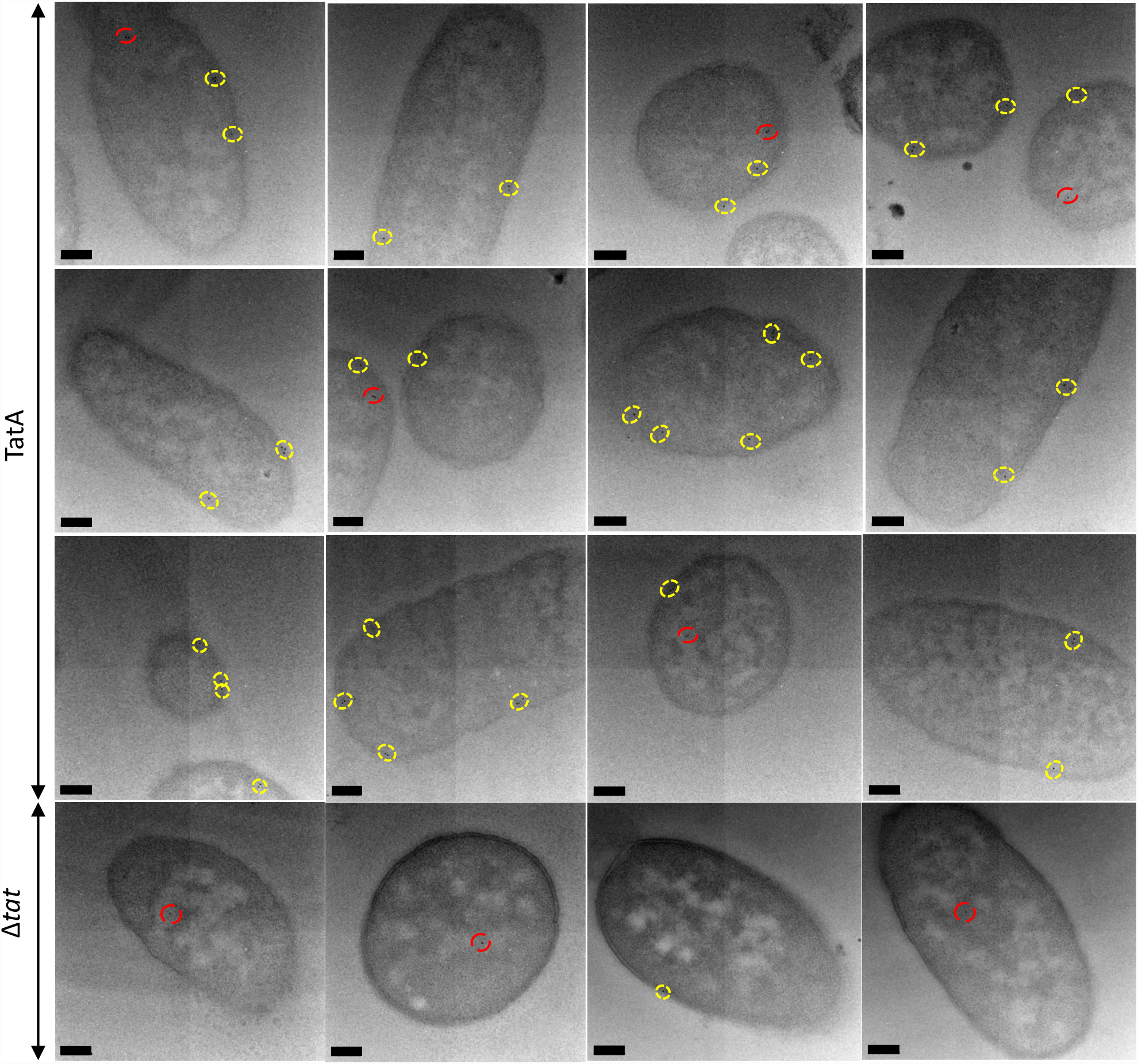
Electron micrographs of *E. coli* cells, overexpressing TatA, immunogold-labelled following primary antibody detection against TatA protein. *E. coli* cells overexpressing TatA, were immunolabelled using a polyclonal antibody raised against TatA (top 3 panels of figure). TatA was found to exhibit a random distribution in inner membrane (yellow circles), and was also present in the cytoplasm (red circles). Controls: cells that did not express Tat machinery, were immunolabelled in the same manner. These cells largely lacked any gold binding, although a few cells bound gold in the cytoplasm and very sparingly at the inner membrane (representative images [labelled (∆*tat*)] in the bottom panel). Images were taken on JEOL-2010F TEM at 12,000X magnification. Scale bar = 200 nm.

Raw gold counts were collected from 200 randomly sampled, immunolabelled *E. coli* TatA-overexpressing and *Δtat* cells, and the gold particles were assigned to either cytoplasmic or inner membrane compartments. Quantification of the gold particles (Figure 3A) shows that the average number of gold particles per compartment in TatA-overexpressing cells was 1.57 per cell (+/− 0.25 gold) in the cytoplasm and 0.99 per cell (+/− 0.2 gold) in the inner membrane, whereas *Δtat* cells bound approximately 0.59 per cell in the cytoplasm (+/− 0.13 gold) and 0.07 (+/− 0.05 gold) in the membrane, respectively. The labelling of the membrane-bound TatA is thus highly specific, with the inner membrane of TatA-overexpressing cells containing 13.2-fold more gold particles than the same membrane in ∆*tat* cells. To confirm a statistical independence in immunogold labelling between these cells at the cytoplasm and inner membrane, chi-squared analysis of the raw gold count data was conducted. For a total chi-squared value of 35.25 and 1 degree of freedom, P was less than 0.005. This confirms that the gold labelling distributions between the two cells types are significantly different.

**Figure 3.**
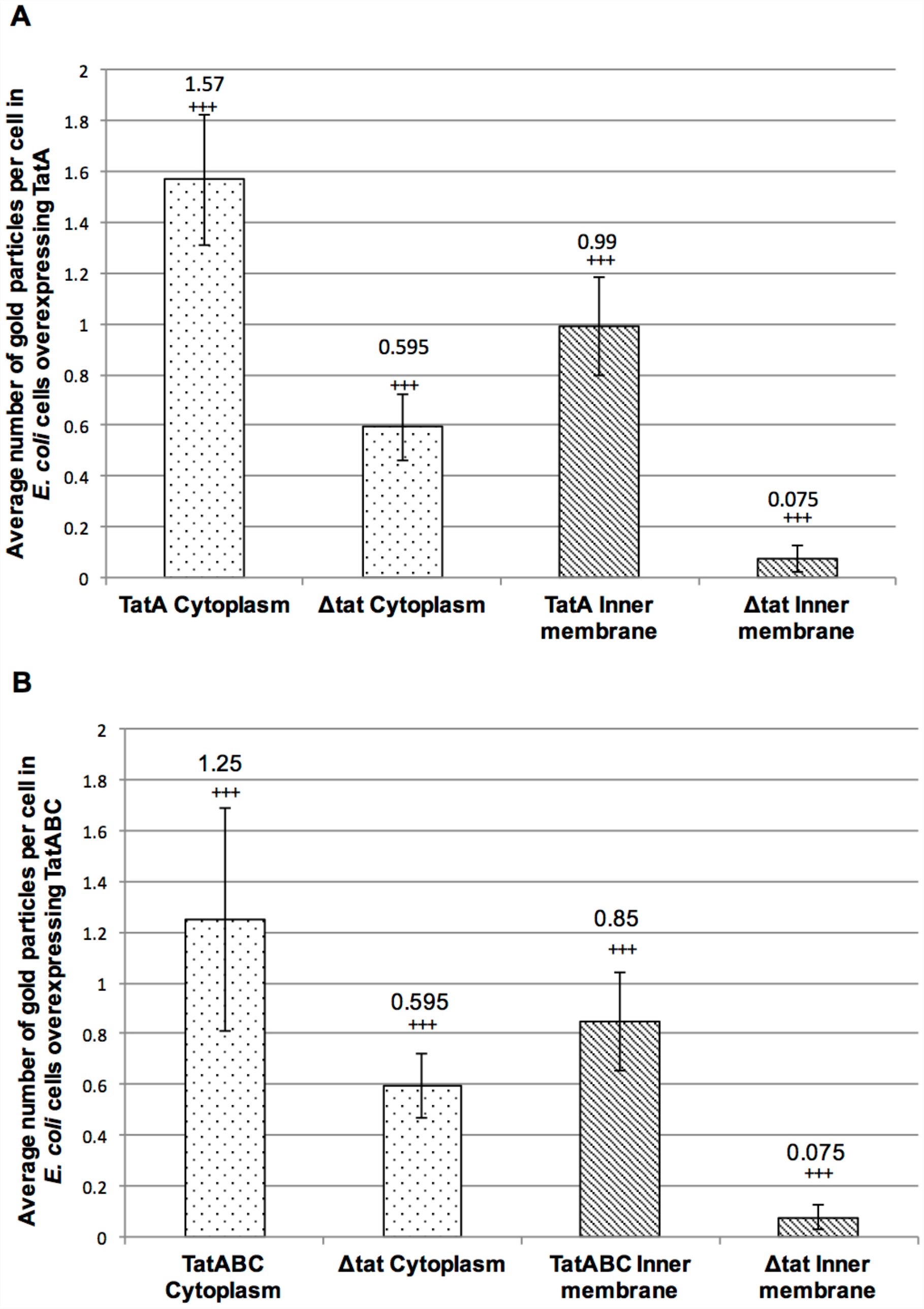
Quantitative analysis of raw gold counts of *E. coli* cells immunogold labelled with primary antibody detection against TatA protein. *E. coli* cells overexpressing TatA or TatABC, were immunolabelled using a primary antibody raised against the TatA protein. Controls were cells that had Tat machinery knocked-out [labelled (∆*tat*)]. Raw gold counts were taken from 200 randomly imaged *E. coli* from 2 separate resin blocks. Gold was assigned to either inner membrane or cytoplasm compartment. The approximate number of gold at the inner membrane and cytoplasm of each cell was calculated for each cell type. A: there were 13.2X more gold at the inner membrane of cells overexpressing TatA than ∆*tat* cells (line-patterned); and 2.64X more gold in the cytoplasm of these cells versus ∆*tat* (dotted pattern). B: there were 11.3X more gold at the inner membrane of cells overexpressing TatA than ∆*tat* cells (line-patterned); and 2.1X more gold in the cytoplasm of these cells versus ∆*tat* (dotted pattern) The control was the same sample in A and B. Error bars: CI of 2xSE. The labelling of TatA between the two cell types is statistically significant (indicated by +++).

Importantly, the gold particles are randomly distributed within the membrane, with no evidence for preferential localisation at the poles or elsewhere. The TatA is overexpressed and in excess over the endogenous TatBC complex, and we therefore conclude that the TatA complex on its own has no intrinsic preference for a specific location in the membrane.

Analysis of the distribution of gold particles in the cytoplasm shows the presence of 1.57 and 0.59 particles in the TatA-overexpressing and *Δtat* cells, respectively. Since the particle number in *Δtat* cells represents nonspecific binding, and assuming that the level of nonspecific binding in wild type cells is the same, this indicates that TatA-overexpressing cells contain an average of 0.97 particles per cell in the cytoplasm, which is about the same as the number in the inner membrane. Some of these particles can be attributed to cytoplasmic TatA, as shown in the blots in Figure 1, but the particle count in the cytoplasm of TatA-expressing cells is nevertheless surprisingly high because the blots showed the majority of TatA to be in the membrane fraction. This higher-than-expected labelling of cytoplasmic TatA relative to membrane-enclosed TatA (in TatA-overexpressing cells) can be explained by the fact that successful antibody binding is reliant on an antigenic site being exposed when sectioning the *E. coli* sample. Given that membranous TatA is enclosed within a phospholipid bilayer, as opposed to the aqueous mileu of the cytoplasm, it is feasible that steric hindrance of the anti-TatA antibody could be imposed by the phospholipid molecules of the inner membrane; resulting in sub-optimal immunolabelling of TatA in the cell wall. This approach thus provides highly specific detection of TatA in the membrane fraction, whereas labelling of cytoplasmic TatA is higher than expected.

### Increased expression of TatBC alters the distribution of TatA in the inner membrane of E. coli

To examine whether the uniform distribution of TatA in the inner membrane is influenced by other constituents of the Tat machinery, we carried out similar analyses of cells that overexpressed the entire TatABC complex rather than only TatA. The cells were immunolabelled with anti-TatA antibody and representative images are shown in Figure 4, with the quantified data shown in Fig. 3B. The data show that TatABC-overexpressing cells bound on average 1.25 gold particles per cell (+/− 0.44 gold) in the cytoplasm and 0.85 particles (+/− 0.2) at the inner membrane, whereas *Δtat* cells bound approximately 0.59 particles per cell (+/− 0.13 gold) in the cytoplasm and 0.07 particles (+/− 0.05 gold) in the membrane. The data thus show that the TatABC cells bind approximately 11.3x more gold at the inner membrane than *Δtat* cells, again confirming that the technique is able to detect membrane-bound TatA with high specificity in these cells. The overall levels of TatA are fairly similar in the 2 types of cell examined: 0.85 particles per cell in TatABC-overexpressing cells and 0.99 in TatA-overexpressing cells, which shows that expression of TatBC does not increase the levels of TatA in the membrane (for example, by stabilising the TatA complexes).

**Figure 4.**
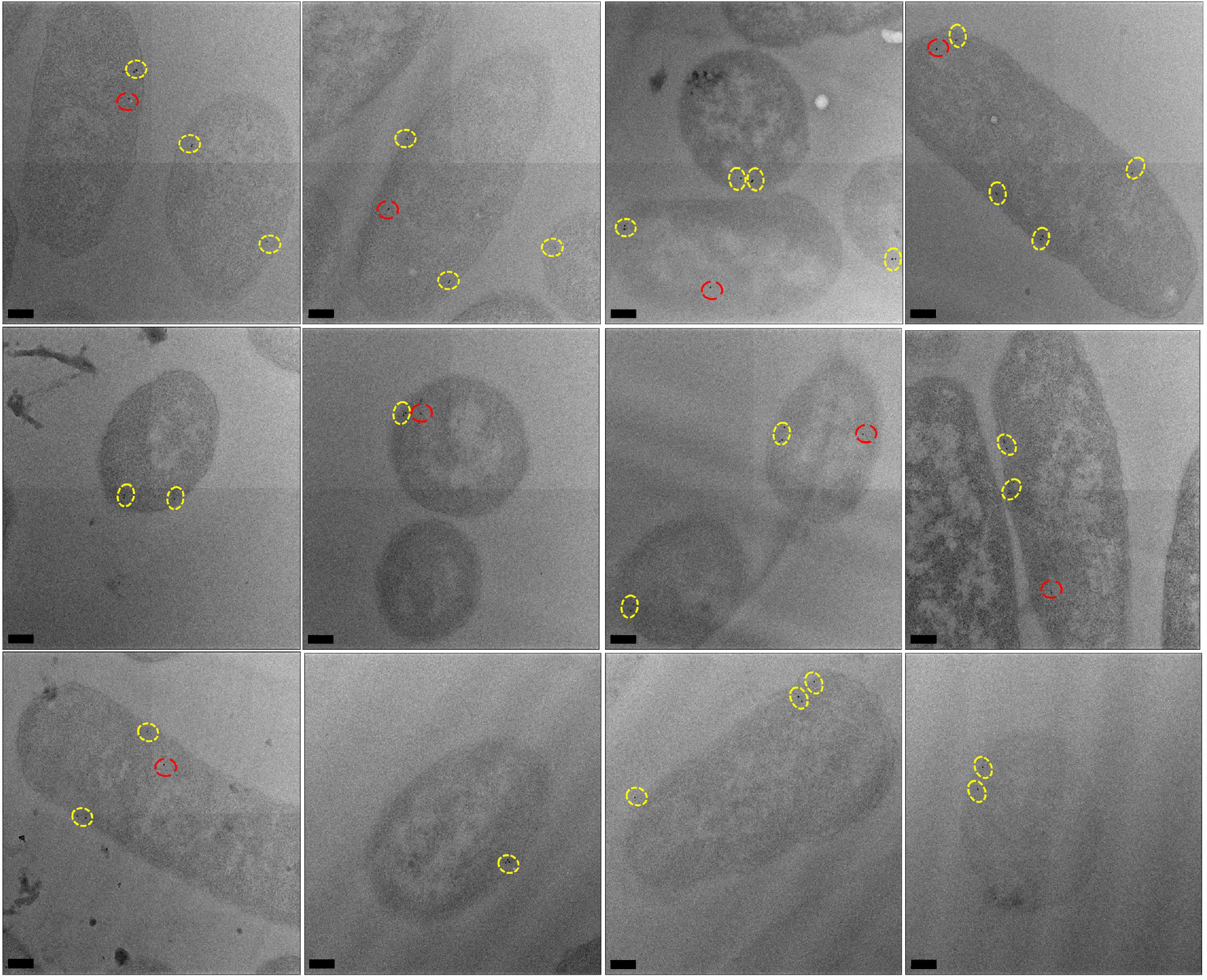
Electron micrographs of *E. coli* cells, overexpressing TatABC, immunogold-labelled following primary antibody detection against TatA protein. *E. coli* cells overexpressing TatABC, were immunolabelled using a primary antibody raised against the TatA protein. TatA protein was found to exhibit a random distribution in inner membrane (yellow circles), and was also present in the cytoplasm (red circles). Images were taken on JEOL-2010F TEM at 12,000X magnification. Scale bar = 200 nm.

Chi-squared analysis confirmed a statistical difference in immunogold labelling between TatABC-overexpressing and *Δtat* cells in both the cytoplasm and inner membrane: for a total chi-squared of 39.2 and 1 degree of freedom, P<0.005. As with TatA-overexpressing cells, there is evidence of non-specific binding of the TatA antibody since TatABC-overexpressing cells bound 1.25 particles per cell in the cytoplasm, while *Δtat* cells bound 0.59 particles. This again contrasts with the relatively low levels of cytoplasmic TatA observed on blots, as discussed in the previous section.

With increased expression of TatBC proteins, immunogold labelling showed TatA to exhibit a uniform distribution around the periphery of most of the independently sampled *E. coli* cells, as shown in Figure 4. However, it was noticed that in a small proportion of cells, the distribution of TatA protein altered in the inner membrane, with linear clustering of gold clearly observed (Figure 5).

**Figure 5.**
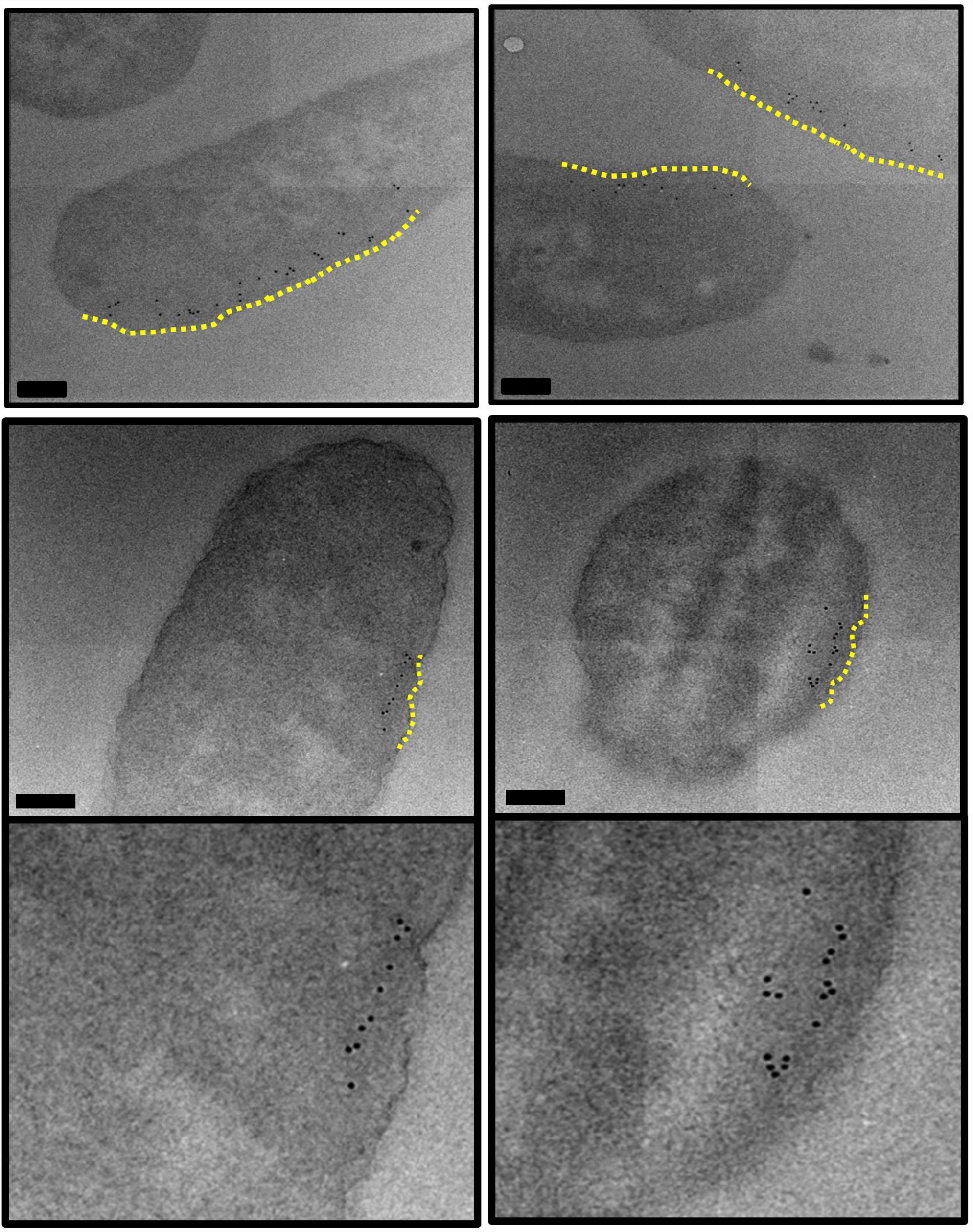
Electron micrographs of *E. coli* cells, overexpressing TatABC, immunogold-labelled following primary antibody detection against TatA protein. *E. coli* cells overexpressing TatABC, were immunolabelled using a primary antibody raised against the TatA protein. With increased expression of TatBC, the localisation of the TatA protein altered in approximately 1 in 25 of cells. TatA protein appeared to cluster along the inner membrane (highlighted in yellow). Images were taken on JEOL-2010F TEM at 12,000X magnification. Scale bar = 200 nm.

Whilst the majority of the gold particles that form these clusters are within 25 nm radius of the cell periphery, a small proportion have bound beyond. It is likely that the high density of antibody labelling in this particular region of the cell (possibly due to the wealth of neighbouring TatA antigenic sites) promotes gold clustering. Nevertheless, all of the gold particles that are classed as being bound to the inner membrane still form a linear clustering along the membrane. This distribution was not observed in any of the TatA-overexpressing cells, and in order to further investigate this previously unseen distribution of TatA in *E. coli*, array tomography was used to visualise an independent sample of the same cell type and gain 3D insight into this very interesting result.

### Array Tomography reveals a linear clustering of TatA protein in the inner membrane of E. coli

Array tomography is a volumetric microscopy technique that involves the physical, serial sectioning of a sample. Unlike traditional serial section TEM approaches (where the section is placed under an electron beam that completely penetrates the specimen), use of a field emission scanning electron microscope (FESEM) to image means that the electron beam is considerable less destructive to the sample, and thus the serial sections are much more stable under the electron beam. This sample stability, in combination with the fact that array tomography can reconstruct considerably larger volumes (since many serial sections can be mounted in the FESEM at one time and imaged in a single session); makes this technique extremely useful in reconstructing large cellular volumes and identifying specific macromolecules *in situ*.

Ultrathin, resin-embedded serial sections of a biological sample are sequentially imaged under the FESEM to obtain 2D backscattered electron micrographs which are then computationally aligned and reconstructed to generate a 3D volume for visualisation. In this study 50 nm serial sections of resin-embedded *E. coli* cells overexpressing TatABC were immunogold labelled (to identify TatA) and subsequently imaged under the FESEM. 3D reconstructions generated insight into the localisation of the gold-labelled TatA protein in 3D space.

Figure 6, top panel, shows a scanning electron micrograph of a single section of immunogold-labelled *E. coli* cells overexpressing TatABC. Figure 6, bottom panel, shows the micrographs obtained from a single cell that was sequentially imaged through 16 separate sections (numbered 1-16; cell outlined in cyan in Figure 6, top panel). Following alignment of the serial sections, the image stack was segmented via manual contouring of the periphery of each individual bacterium, on each of the sections. Simultaneously, the positions of any 10 nm gold particles present on the section was also marked (lower panel, red circles). Gold particles were clearly distinguished from image noise and contaminants in the serial sections as spherical bodies, 10 nm in diameter. This procedure was repeated for 6 cells chosen at random.

**Figure 6.**
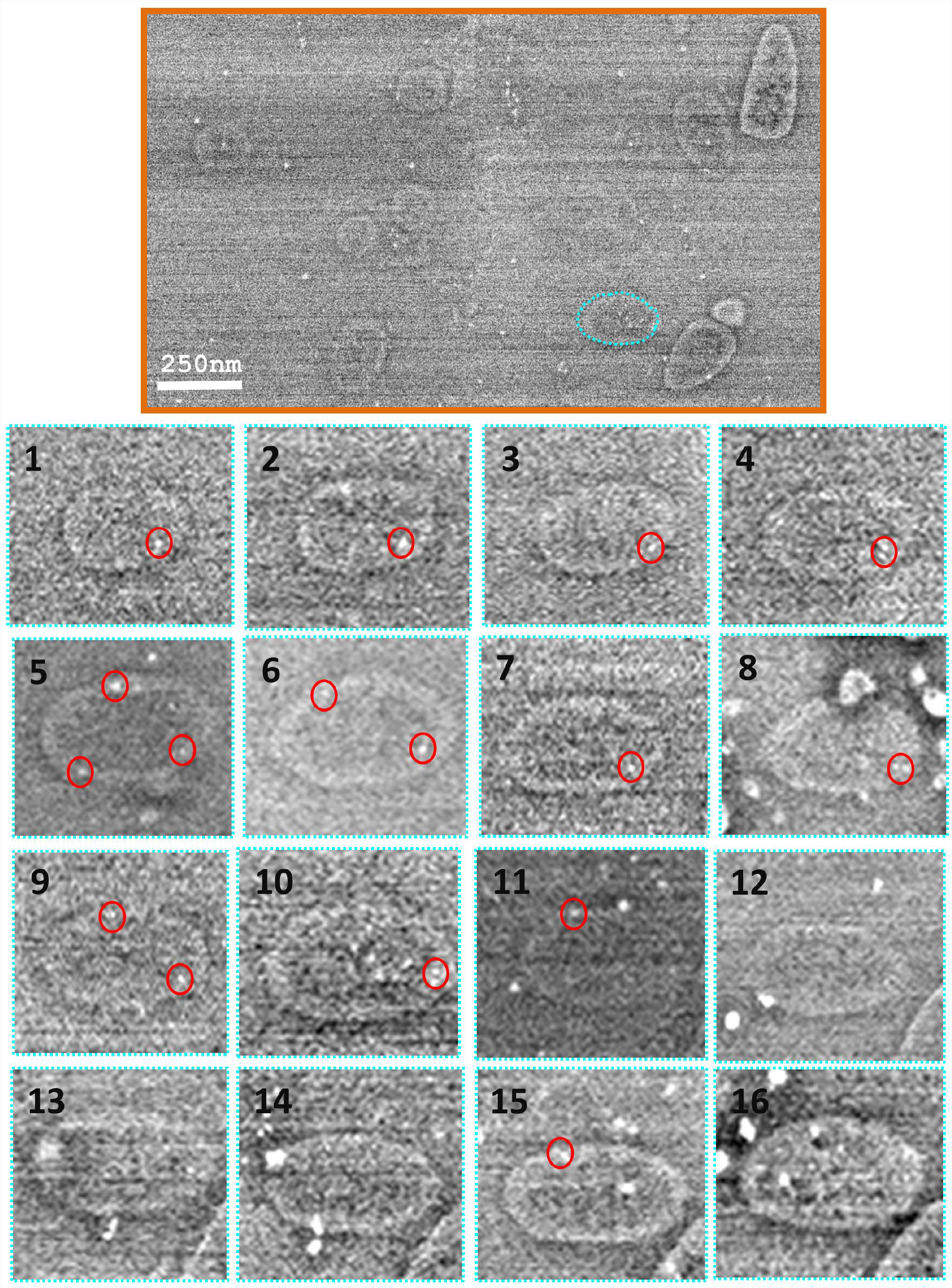
Alignment and segmentation of array data. Top panel: Having sequentially imaged the serial sections (of resin-embedded *E. coli* cells overexpressing TatABC) at 5000X magnification, the data was aligned and segmented for generation of a 3D model. Bottom panel: Using IMOD software, individual *E. coli* cells were manually contoured via tracing the entire perimeter of the cell on each section (images 1-16 below). Also, the position of any gold (easily identified as a spherical 10 nm contrast under the electron beam) was added to the model (red circles below on images 1-16). IMOD software was used to mesh sequential contours for generation of 3D reconstructions of *E. coli*. Images taken on Image taken on JEOL-7401F. Scale bar = 250 nm.

Having successfully contoured the *E. coli* cells that were visible in all of the 20 sections, 3D reconstructions of individual bacteria were generated (Figure 7). Analysis of the positions of the gold particles (and thus TatA protein) revealed a linear cluster of TatA in each *E. coli* cell-shown in Figure 7. Measuring the distance of these gold particles in the 2D micrographs from the cell periphery revealed that they are all located within 25 nm of the inner membrane and could thus be classed as localised to this cellular compartment. Furthermore, the clusters contained up to 10 TatA-bound gold particles; suggesting that TatA formed clusters in the inner membrane up to 0.5 µm in length (based on the knowledge that the gold particles are located at 50 nm intervals owing to the thickness of an individual section).

**Figure 7.**
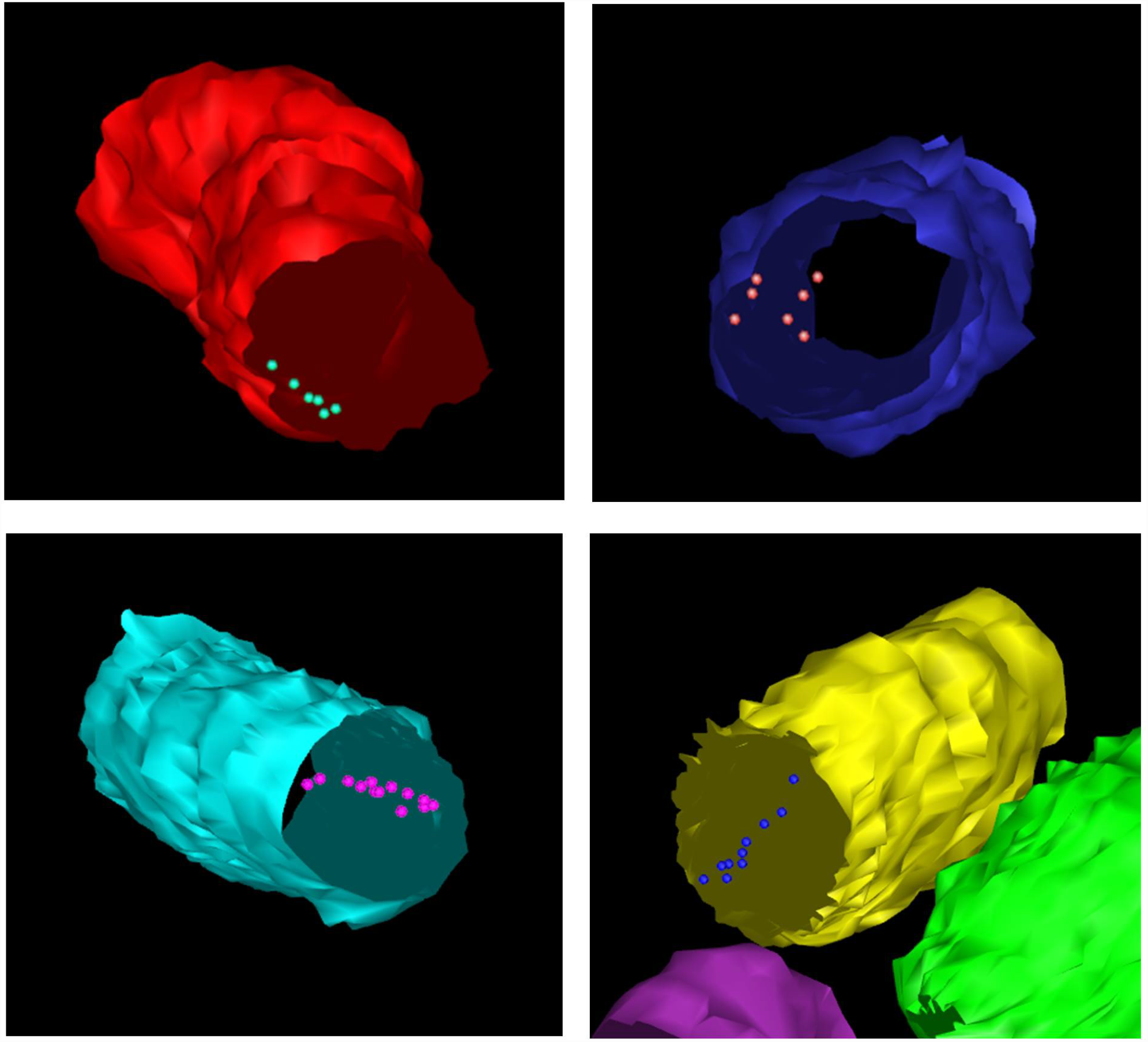
3D visualisation of a linear distribution of the TatA protein in the inner membrane of *E. coli*. Upon reconstruction of individual *E. coli* cells, the position(s) of immunolabelled TatA protein could be deciphered. The above reconstructions illustrate a linear clustering of TatA protein in the inner membrane of *E. coli*.

The array tomography reveals linear clusters which would not have been observed without alignment of these 2D sections. This suggests that the linear clusters are much more common than was suggested by our earlier 2D analysis in the previous section of this manuscript. This is the first time that the localisation and distribution of any Tat component has been visualised using 3D approaches, and the data suggest that such approaches may provide valuable insights into Tat complex organisation.

## Discussion

TatA plays an essential role in the Tat translocation process, and insight into its localisation and distribution in cells is fundamental to elucidating the Tat translocation pathway. Analysis of this membrane protein *in situ*, avoiding the use of detergents, remains a challenging task. Previous experiments have used YFP-labelled TatA proteins to analyse Tat distribution in the inner membrane (Leake et al., 2008, Alcock et al., 2013). However, TatA-XFP fusion proteins are not fully functional(Ray et al., 2005, Alcock et al., 2013). In this study we have used 2D and 3D EM approaches to study the location and organisation of TatA which is native, apart from the presence of a C-terminal His tag which has been shown not to affect activity(Bolhuis et al., 2001). This is thus the first microscopic study to analyse native TatA complexes and the first to analyse TatA organisation in 3D. Given that this translocase accomplishes the remarkable feat of transporting large, fully-folded proteins without jeopardising membrane integrity, such spatial data are important for an understanding of the system.

The 2D immunogold labelling of TatA-overexpressing cells shows that the protein is randomly located within the plasma membrane when present in excess over the TatBC proteins. There is no evidence for preferential location at the cell poles or elsewhere, and we observed no evidence for any supermolecular organisation of TatA complexes on their own. The labelling of the TatA in the membrane is highly specific and on the basis of these results it appears that TatA complexes are able to adopt random positions in the plasma membrane at steady state, when the levels of overexpression are such that most of the complexes cannot be engaged in translocation due to lack of TatBC complexes.

Whilst overexpression of TatA protein makes it easier to detect during microscopy analysis, it could be argued that this increased expression could potentially result in protein mislocalisation. However, overexpression of Tat machinery is necessary owing to the low expression levels of native Tat machinery in bacteria. As expected, immunogold labelling of *E. coli* expressing wild-type levels of Tat machinery failed to bind any anti-TatA antibody and thus no gold was seen in any of the samples (data not shown). Furthermore, it is known that overexpression of Tat machinery is not detrimental to the cell, and overexpression of TatABC is able to relieve saturation of wild-type Tat machinery when substrate is overexpressed, showing that overexpressed Tat complexes are biologically active (Matos et al., 2012).

In contrast to the above, the 2D immunogold labelling of cells that concomitantly overexpress the TatBC proteins provides the first signs of an interesting change in organisation in Tat machinery in response to alterations in Tat subunit stoichiometry. Using 2D TEM, a relatively small proportion (4%) of the cells could be seen to exhibit a change in localisation of TatA. However, for such clusters to be observed in individual 2D sections, they must be perfectly aligned with the direction of the cut. This makes it highly likely that many clusters in other cells were missed due to the sectioning process. When using the 3D approach of array tomography all linear clusters within the volume analysed would be visible.

3D array tomography was able to provide a more detailed analysis of TatA localisation. It was shown that the TatA protein of the twin arginine translocase forms linear clusters in the inner membrane of *E. coli* in a TatBC-dependent manner. This is not a strictly linear ‘tubule’ type of pattern but rather an extended cluster of particles. Although the sample size is small, this is the first demonstration of such an organisation for the TatA complex. Previous studies on TatA overexpression(Berthelmann et al., 2008) observed relatively massive TatA tubules in *E. coli*, but these were located in the cytoplasm and those tubules seem entirely different to the clusters observed here. However, we recently showed that a mutation in the TatAd subunit of the *Bacillus subtilis* TatAdCd system triggered the formation of long fibrils when expressed in *E. coli*, and we speculated that this could be an aberrant form of super-assembly (Patel et al., 2014). It is possible that the clusters observed in the present study are related to those fibrils: it could be that the TatA subunit of the Tat translocase machinery has the propensity to form long oligomeric assemblies inside both Gram-positive and Gram-negative bacteria.

Other studies have used XFP-labelled TatA to study its distribution and assembly using optical techniques, but the major problem in these cases is the finding that TatA-XFP on its own is inactive (Ray et al., 2005, Alcock et al., 2013). Alcock *et al.* (2013) expressed a TatA-YFP fusion in the presence of TatE, a TatA paralogue that is capable of functionally replacing TatA, and observed a doubling of export of a native Tat substrate. They observed that the TatA can exist in both dispersed and higher oligomer states and that the distribution of TatA complexes is controlled by the other components of the Tat machinery, namely TatBC. However, it is impossible to properly reconcile those data with the EM study reported here for two reasons. First, the Alcock *et al*. study may have actually measured assembly-disassembly of TatE complexes, with molecules of inactive TatA-YFP incorporated into the TatE complexes, and secondly, it is unclear how TatA-YFP complexes lacking TatE may have contributed to the fluorescence signals.

To conclude, the novel combinatorial approach of immunogold labelling of *E. coli* cells with array tomography has enabled us to gain new insight into the 3D localisation and distribution of Tat complexes in their native cellular environment. Array tomography has identified a previously uncharacterised distribution of TatA, however, at present we do not know whether it is consistent with a pore-forming mechanism of Tat. Further immunogold co-labelling studies could potentially provide additional insight into the relative location of Tat proteins in the inner membrane, however the immunolabelling technique is reliant on using an antibody of very high affinity. We attempted immunolabelling experiments using anti-TatB and anti-TatC antibodies, but sufficiently specific labelling with these antibodies could not be demonstrated and failed to label the *E. coli* sections.

Interestingly, confocal microscopy experiments have revealed that the Sec translocase has the ability to form large helical assemblies inside of *E. coli* (Shiomi et al., 2006), and it is thus possible that the linear clustering of TatA witnessed in this study is the first indication of a similar higher-order assembly of the Tat machinery in bacteria. The true significance of such higher-order assemblies remains unclear, but the techniques outlines in this study have significant potential for addressing the outstanding questions in this area.

## Materials and Methods

### Preparation of E. coli cells for visualisation by electron microscopy

Chemical fixation of *E. coli* cells: *E. coli* cells (overexpressing TatA or TatABC, on Δtat background) were resuspended in aldehyde fixative (0.25% glutaraldehyde/4% formaldehyde) and low melting temperature agarose at 1:1 volume ratio. Agarose-enrobed cells were then harvested and incubated on ice until the agarose had set. Cells were diced into 1 mm^3^ pieces using a clean razor blade and resuspended in fresh aldehyde fixative for overnight fixation at 4 °C. Chemically fixed cells were washed for 1-2 hrs in 0.15 M sodium cacodylate/HCl buffer pH 7.4 at 4 °C to remove aldehyde fixative. The reaction was then quenched for 1hr by washing in cacodylate buffer containing 0.1M glycine. Contrast was imparted to the cells by incubation in 1% tannic acid (TA) for 1 hr at 4°C, followed by washing in H_2_O for a second staining of 1% uranyl acetate (aq) for 1 hr at 4°C. *E. coli* cells were dehydrated in an ethanol series (70-100%) for a period of 2hrs for subsequent resin embedding (London Resin Company Ltd.; Berkshire, UK).

### Serial sectioning of resin blocks for imaging by electron microscopy

Ultrathin (50 nm) serial sections were cut (using an ultramicrotome [Ultracut E, Reichert-Jung; Vienna, Austria]) and transferred to either carbon-coated (~80 nm thickness) glass coverslips (13 mm diameter) for scanning EM work, or 200-mesh carbon-coated copper grids (Agar Scientific, Stansted Essex) for transmission EM work. Serial sections were blocked via incubation in TBS/Tween buffer (0.24% Tris (w/v), 0.8% NaCl (w/v) and 0.1% Tween-20 (v/v) pH 8.4) containing 1% BSA (w/v) and 4% normal goat sera (v/v) (Abcam Plc, Cambridge UK) for 30 mins at room temperature (RT). TatA protein was labelled by incubation with primary anti-TatA antibody (rabbit monoclonal, as described in (Bolhuis et al., 2000)) at 1:20 dilution (TBS/T +1% BSA buffer) for 2hrs at RT. Sections were then washed with TBS/T +1% BSA at RT, and then incubated with 10nm gold-conjugated secondary antibody (Goat anti-rabbit IgG pre-adsorbed, Abcam Plc) for 1 hr at RT. Finally, sections were washed in TBS/T (+1% BSA) and dH_2_O and allowed to air dry before insertion into the EM.

### Fractionation of E. coli cells and detection of TatA protein

*E. coli* cells were fractionated into periplasmic, cytoplasmic and membrane fractions using the lysozyme/cold osmotic shock method (Randall, 1986). The resulting fractions were analysed by sodium dodecyl sulphate polyacrylamide gel electrophoresis (SDS PAGE).

Once the cytosolic and inner membrane samples had been analysed using SDS-PAGE the proteins were transferred from acrylamide gels to PVDF membranes via semi-dry Western Blotting apparatus. The membranes were blocked overnight in a solution of 5% (w/v) dried milk in PBS-T and then incubated with primary anti-TatA antibody (1:6000, in PBS-T) for 1 hour before washing. Membranes were incubated with anti-rabbit-HRP conjugate (Promega, WI, USA) for 1hr at RT, and subsequently washed. Finally, immunoreactive bands were detected using ECL^TM^ detection reagents according to the manufacturer's instructions. X-ray films were developed using an AGFA Curix 60 automatic developer as directed by the manufacturer's instructions.

### Quantification of immunogold labelling

A multi-random sampling approach was used to analyse the distribution of gold within the immunolabelled *E. coli* sections. *E. coli* specimens were sampled in a hierarchical fashion, commencing at the liquid culture stage: ¼ of the culture media was randomly selected and harvested for subsequent downstream processing (Fig S1). Specimens were blinded at the point immunolabelling, such that the distribution of gold was analysed objectively. Raw gold counts from 200 randomly imaged *E. coli* (from 2 biological and 2 technical replicates) were recorded as numerical frequency distributions and assigned to one of the two compartments: inner membrane and cytoplasm. Raw gold counts were used to construct a numerical frequency distribution table. Observed numbers of gold particles in each compartment were compared by use of contingency table analysis with ‘g’ groups (arranged in columns) and ‘c’ compartments (arranged in rows). This analysis generated predicted gold particles and, hence, partial chi-squared values, for each group and each compartment. Comparison of the total chi-squared value against a Chi-squared table determined whether the null hypothesis (of no difference in immunolabelling between TatA overexpressing cells and Δtat cells) could be rejected (p<0.005).

### Analysis of arrays to generate volume image of E. coli bacteria

Serial sections were sequentially imaged using a JEOL-7401F field emission scanning electron microscope at 5 kV accelerating voltage and 5000X magnification. SEMSupporter software (System in Frontier Inc.) enabled the acquisition of image montages from each section for generation of an image stack.

### Alignment of image stack

The Panorama Mapping module in SiFi SEM Supporter software was used to manually refine alignment of the montage stack. The greyscale of the images was reset for each montage to make refinement of alignment easier in later steps. In eTomo software(Mastronarde, 1996), Align Serial Sections/Blend Montages tab was selected and the relevant .mrc image stack uploaded. Midas module was used to further manually refine montage alignment; correcting for magnification, stretch and rotation. Under the ‘Make Stack’ tab in Align Serial Sections, Global Alignments was selected, and Make Aligned Stack option selected to generate aligned volume stack.

### Segmentation of aligned image stack for generation of volume image

Following alignment using Midas, the Z-stack was opened using 3DMod (set to Model). The first Z-plane was selected, and the first bacterium centred, ensuring it filled the screen. A new Object was created for each bacterium.

For each Z-plane (1 to 20), the perimeter of a single *E. coli* bacterium was manually traced. Any immunogold labels were each marked as single points in each contour. Points were edited and deleted as necessary to make sure that the contour matched the visible perimeter as accurately as possible. Once the contour was complete, any immunogold labels were each marked in a new Object as separate contours. The perimeter tracing and gold-point marking was repeated for each Z-plane until all were successfully contoured.

The contours were stitched together using the meshing feature in 3DMod. Mesh One was used to generate the surface for each bacterium. The final model was saved and exported into UCSF Chimera for visualisation.

## Acknowledgements

We would like to thank Ian Hands-Portman (Imaging suite, University of Warwick) and Dr Kirsty MacLellan-Gibson (Imaging Sciences, National Institute for Biological Standards and Control) for allowing continued access to TEM and SEM machines, respectively. We are grateful to the Welcome Trust for generous support (Grant 055663/Z/98/Z) to the Imaging Suite at the University of Warwick.

### Competing Interests

The authors declare no competing interests

### Author contributions

SMS performed all of the experiments, analysed the data and wrote the manuscript. AY assisted in the use of the scanning electron microscope during image acquisitions, and provided assistance in the analysis of the EM data to generate a 3D volume. CJS and CR helped to write the manuscript. CJS and RF conceived the project.

### Funding

This work was funded by the Biotechnology and Biological Sciences Research Council (BBSRC) and the Wellcome Trust (055663/Z/98/Z).

---

**Figure S1.**
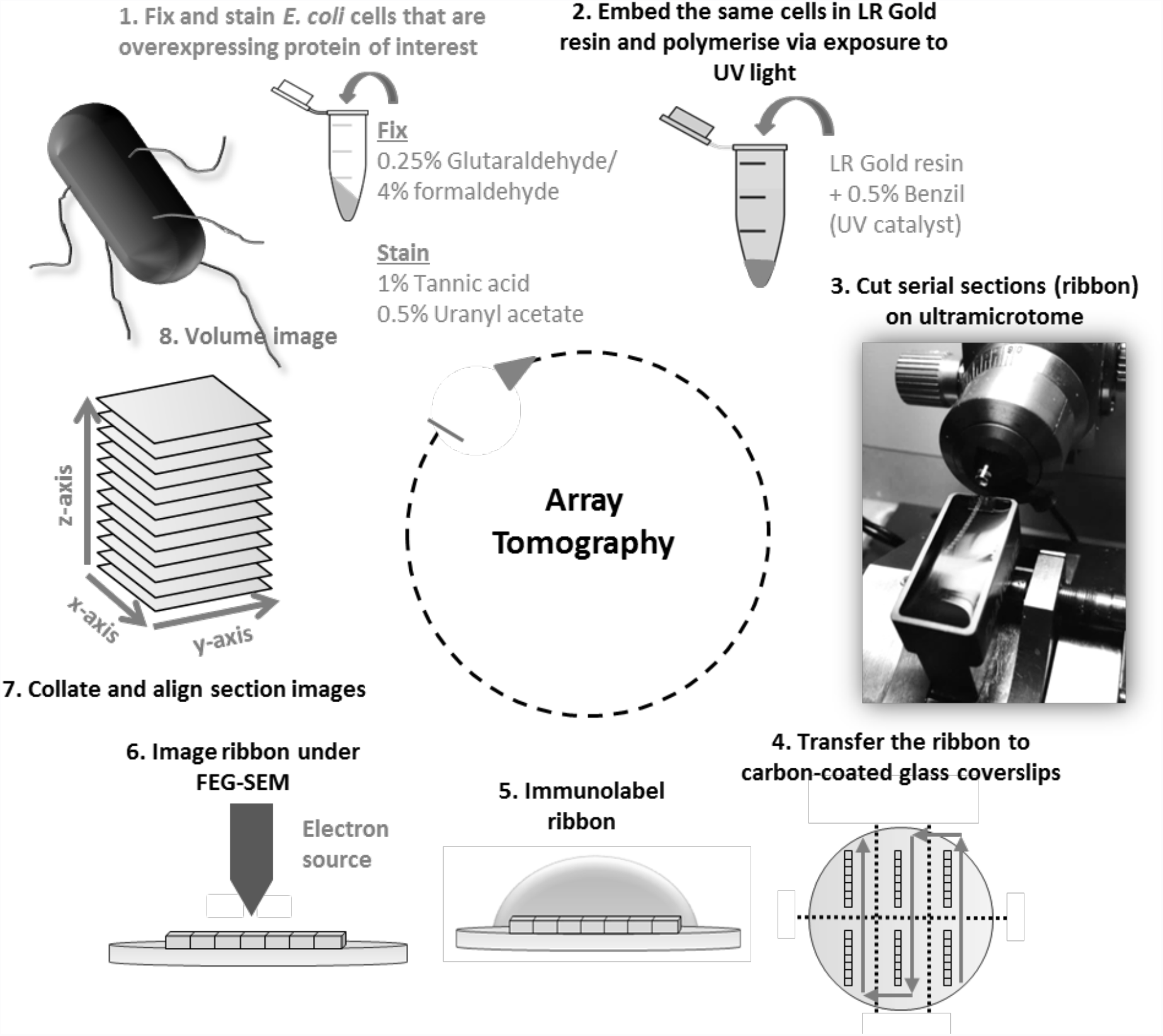
Overview of the Array Tomography process. Array tomography is an 8-step technique that results in a volume image of the sample studied, in this case an *E. coli* bacterium. Fixation was necessary to biologically preserve the *E. coli* cells, and staining generated contrast under the electron beam (Step 1). Embedding in a resin forms a solid medium that enables the sample to be imaged in the microscope (Step 2). This requires the *E. coli* bacteria to be cut using a diamond knife (Step 3). Ultrathin (50 nm) serial sections were transferred to a flat, conductive surface for visualisation under the scanning electron microscope (SEM) (Step 4). The protein of interest shall be identified by exploiting the presence of a specific antigen i.e. TatA protein (Step 5). Imaging the ribbon under the FESEM (Strep 6) generates an image stack (Step 7) that is to be computationally rendered to generate a 3D reconstruction of an *E. coli* bacterium (Step 8).

**Figure S2.**
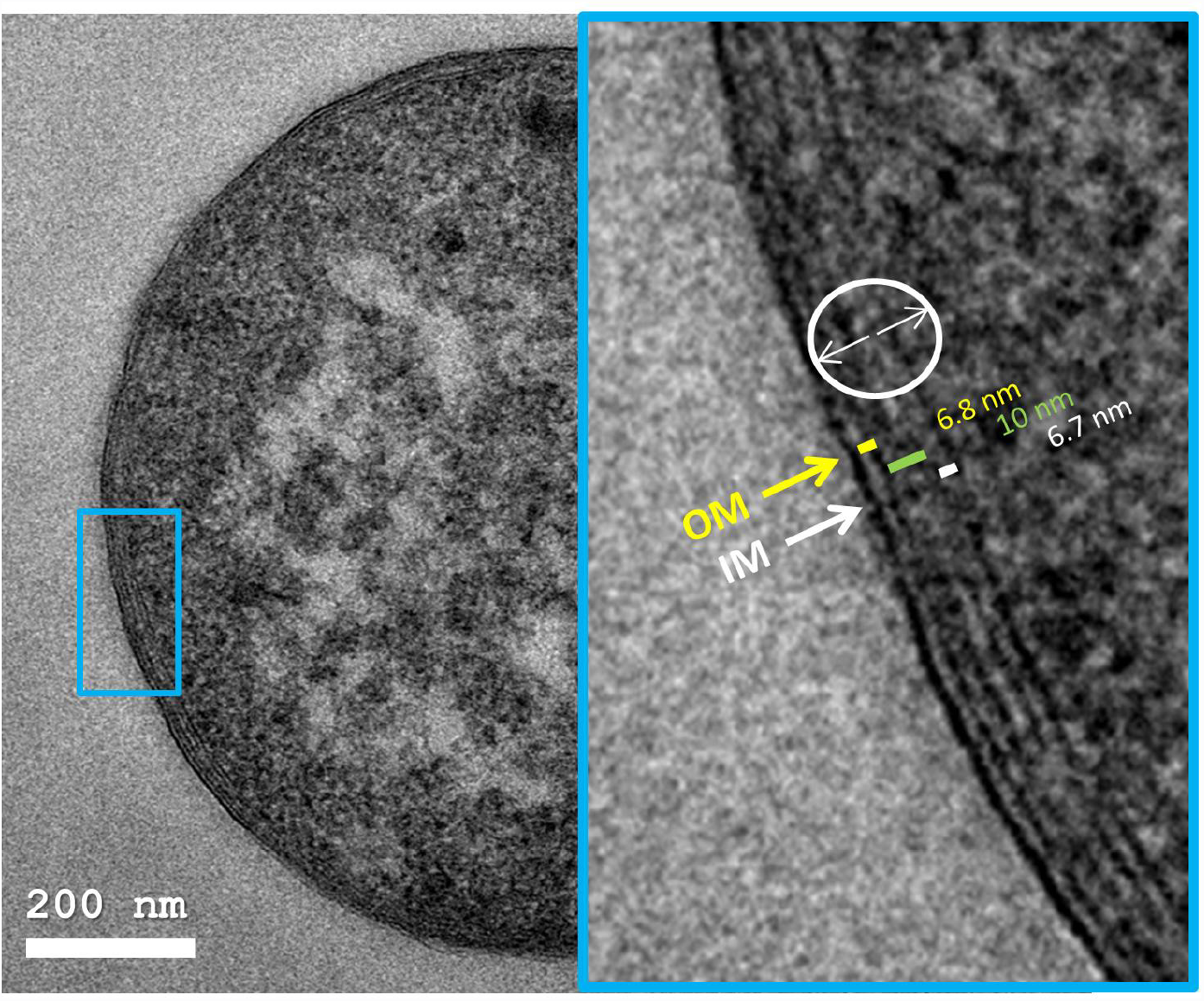
Electron micrograph of a sectioned *E. coli* bacterium. A: Electron micrograph of an *E. coli* cell stained with tannic acid and uranyl acetate. The *E. coli* bacterium shown has been sectioned through latitudinal axis with a diamond knife. The aforementioned heavy metal staining procedure yields good internal contrast such that the cytoplasmic and cell wall features are clearly distinguishable. B: Zoomed image of *E. coli* cell wall from A. The measurements of the outer membrane (OM, yellow labelling), periplasm (green labelling) and inner membrane (IM, white labelling) are consistent with average measurements for these cellular compartments of *E. coli*. Gold particles within 25 nm radius of the section shown (white circle) shall be considered as bound to the inner membrane of *E. coli*. Image taken on JEOL-2010F TEM at 5000X mag. Scale bar = 200 nm.

**Figure S3.**
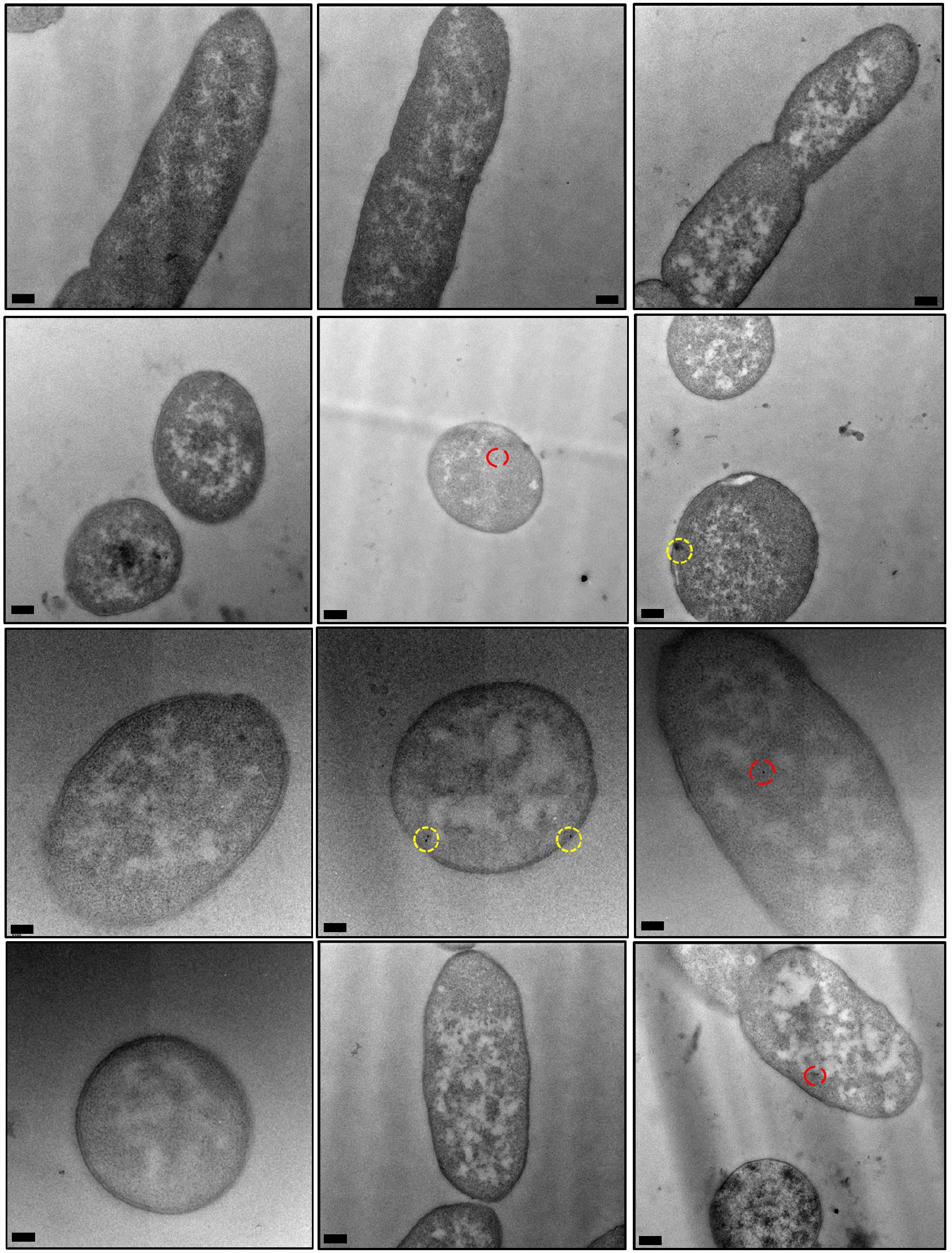
Electron micrographs of *E. coli* cells, lacking expression of Tat machinery, immunogold-labelled following primary antibody detection against TatA protein. *E. coli* cells lacking expression of any Tat machinery were immunolabelled using a primary antibody raised against the TatA protein to serve as a negative control. An abundance of cells lacking gold binding were detected. A minority of cells bound gold at the cytoplasm (red circles) and the inner membrane (yellow circles). However, the difference between these control cells and cells overexpressing TatA protein was confirmed to be statistically significant. Images were taken on JEOL-2010F TEM at 12,000X magnification. Scale bar = 200 nm.

**Figure S4.**
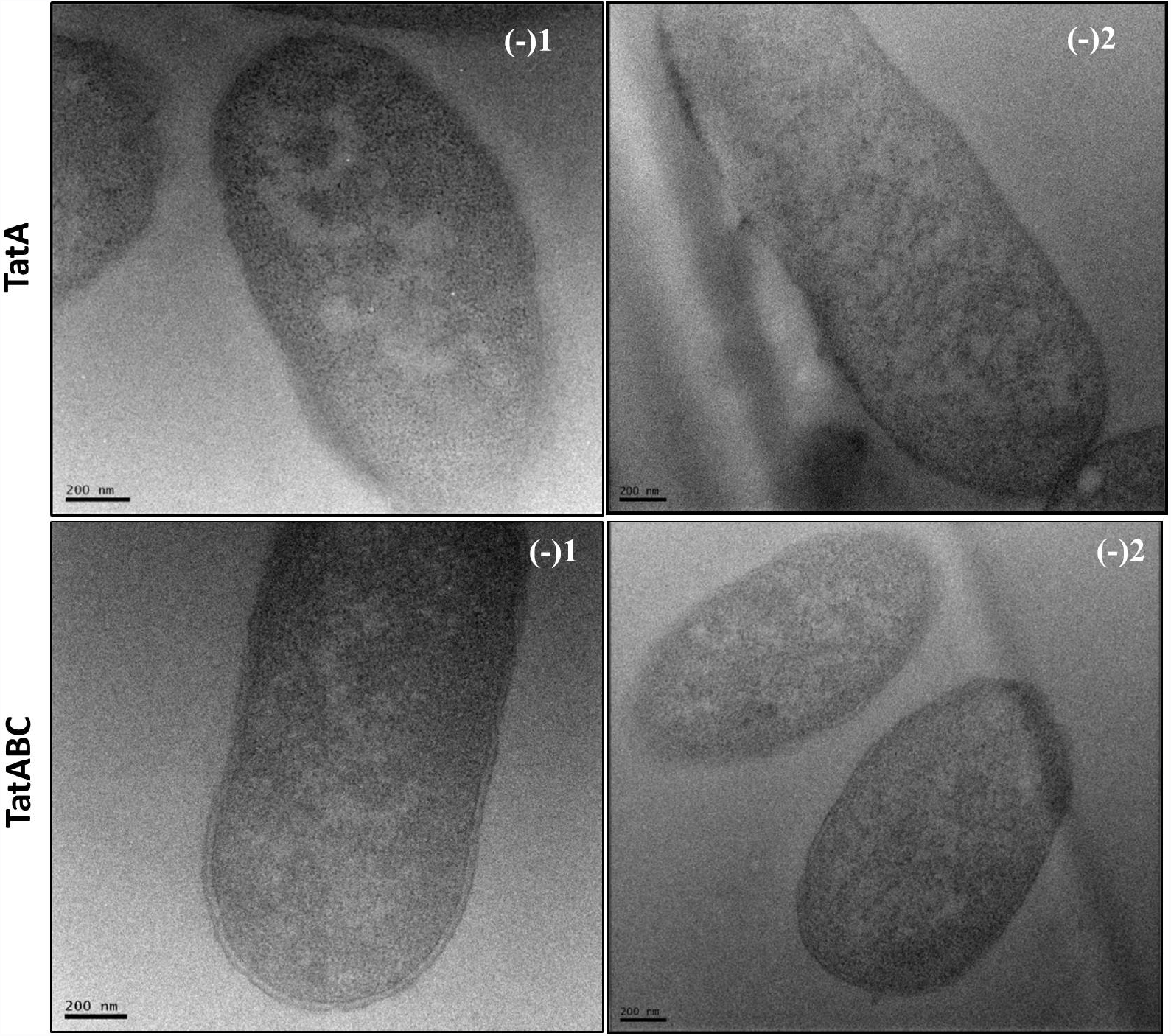
Electron micrographs of *E. coli* cells, overexpressing TatA or TatABC, immunogold labelled omitting the use of primary antibody and using a different secondary antibody. *E. coli* cells overexpressing TatA (top panel) or TatABC (bottom panel) had the primary antibody omitted [labelled (-1)], and also immunolabelled using a different secondary antibody [labelled (-2)]. These cells lacked any gold binding, thus confirming that the secondary antibody does not bind non-specifically to the sample.

